# A post-inflammatory C3-high astrocyte state persists after inflammatory stimulus withdrawal and is attenuated by JAK inhibition

**DOI:** 10.64898/2026.05.26.725945

**Authors:** Yogo Sakakibara, Kyohei Okahara, Jungo Kakuta, Kazumi Emoto, Yuri Ohfusa, Kenji Ohba

**Author notes:** These authors contributed equally to this work (Yogo Sakakibara, Kyohei Okahara). Correspondence: Yogo Sakakibara (Primary Corresponding Author), Kenji Ohba (Co-Corresponding Author).

## Abstract

Reactive astrocytes contribute to neuroinflammation and synaptic dysfunction, but it remains unclear whether transient inflammatory stimulation causes a persistent reactive state after the initial inflammatory stimulus is removed. Here, we investigated whether transient exposure to a defined inflammatory cytokine/complement cocktail induces a persistent reactive astrocyte state and examined the signaling mechanism underlying its maintenance. Human astrocytes were exposed to the inflammatory stimulus and subsequently subjected to stimulus washout, followed by time-course analyses to compare the reversibility of inflammatory gene expression after stimulus removal. Following washout, the expression of several inflammatory response genes, including CXCL10 and NF-κB-associated genes such as NFKBIA, TNFAIP3, and RELB, returned toward baseline levels. In contrast, C3 expression remained elevated, indicating persistence of a post-inflammatory C3-high astrocyte state after withdrawal of the inflammatory stimulus. Pharmacological inhibition of JAK signaling reduced persistent C3 expression to near-baseline levels, supporting the involvement of JAK-dependent signaling in maintenance of this persistent state. Together, these findings suggest that transient inflammatory stimulation induces a post-inflammatory persistent C3-high astrocyte state that is maintained even after broader inflammatory gene responses have subsided. This persistent C3-high component is pharmacologically attenuated by JAK inhibition, identifying JAK-dependent pathways as modulators of persistent astrocyte inflammatory reactivity.

## BACKGROUND

Neuroinflammation is increasingly recognized as a key component of many neurological disorders, including Alzheimer’s disease, multiple sclerosis, amyotrophic lateral sclerosis, and other neurodegenerative or neuroinflammatory conditions [1, 2]. Although inflammatory responses can contribute to host defense and tissue repair in the acute phase, chronic or dysregulated neuroinflammation is thought to promote synaptic dysfunction, neuronal injury, and disease progression [1, 3]. Astrocytes are not merely passive responders to inflammatory cues but active participants in neuroinflammatory signaling [4, 5]. Through bidirectional interactions with neurons, microglia, vascular cells, and infiltrating immune cells, reactive astrocytes help shape local inflammatory environments and influence CNS pathology [4–6].

Under physiological conditions, astrocytes contribute to central nervous system homeostasis by supporting synaptic activity, regulating extracellular ions and neurotransmitters, maintaining metabolic support, and contributing to blood–brain barrier function [7–10]. In diseased or injured brain, however, astrocytes undergo context-dependent transitions into reactive states [11, 12]. Reactive astrocytes comprise heterogeneous and dynamic cell states with distinct molecular, morphological, and functional properties, rather than a single uniform phenotype [11–13]. In inflammatory settings, reactive astrocytes can produce cytokines, chemokines, and complement-related molecules that shape local neuroinflammatory responses and influence neuronal and synaptic integrity [11–13].

Activated microglia have been shown to induce a neuroinflammatory astrocyte program through secretion of interleukin-1α (IL-1α), tumor necrosis factor-α (TNF-α), and complement component 1q (C1q) [12]. This cytokine combination has been widely used as an experimental paradigm to induce inflammatory reactive astrocyte features, often referred to as A1 induction [12]. While the A1/A2 terminology remains useful for describing experimentally induced inflammatory astrocyte features, reactive astrocyte states are increasingly understood to be diverse and context dependent [5]. Among molecules associated with this microglia-induced inflammatory astrocyte program, complement component 3 (C3) is one of the most widely used readouts because it is robustly induced in microglia-triggered reactive astrocytes and has been implicated in complement-mediated synaptic pathology [12, 14, 15].

Because inflammatory astrocyte states have been implicated in neurodegenerative and neuroinflammatory pathology, pathways that regulate astrocyte reactivity or its downstream consequences have been considered potential therapeutic targets [12, 15–17]. These include blocking the induction of inflammatory astrocyte states, reducing astrocyte-derived inflammatory or complement-related signals, and targeting intracellular pathways involved in reactive-state maintenance [12, 15–17]. However, it remains unclear whether inflammatory astrocyte states can be mitigated by simply removing the inflammatory stimulus. Although recent work has begun to examine the reversibility of human reactive astrocyte states after cytokine withdrawal, most studies have focused on the induction phase or established reactive astrocytes, and the persistence and maintenance mechanisms of specific post-inflammatory astrocyte features remain less characterized [18]. This may be relevant to chronic neurological diseases, in which inflammatory stimuli may fluctuate over time, yet astrocyte-driven pathological features may continue to influence neighboring glia and synapses even after the initial inflammatory stimulus has been diminished. Therefore, identifying the pathways that maintain post-inflammatory astrocyte states could provide insight into therapeutic strategies for controlling sustained neuroinflammatory responses.

In this study, we investigated whether transient exposure of astrocytes to A1-inducing inflammatory stimulation leaves a persistent reactive astrocyte state after stimulus withdrawal. We found that although many inflammatory-response genes returned toward baseline after withdrawal of the inflammatory stimulus, C3 expression remained elevated, indicating a post-inflammatory persistent C3-high astrocyte state. This state was suppressed by Janus kinase (JAK) inhibition, suggesting a role for JAK-dependent signaling in its maintenance. Together, these findings support a model in which transient inflammatory stimulation can leave a persistent but pharmacologically modifiable astrocyte state after removal of the initiating stimulus.

## METHODS

### Human primary astrocyte culture and inflammatory stimulation

Human primary astrocytes were obtained from ScienCell Research Laboratories (Cat. No. 1800-10) and used across two independent product lots (lot nos. 35416 and 29851). Unless otherwise indicated, experiments shown in the main figures were performed using lot no. 35416. RNA-seq analysis included samples from both lots, and key findings related to persistent C3 expression and JAK-inhibitor-mediated attenuation were further confirmed using lot no. 29851.

According to the manufacturer, cells were cryopreserved at passage 1 and delivered frozen. Upon receipt, cells were thawed, expanded once in Astrocyte Medium (ScienCell Research Laboratories, Cat. No. 1801) supplemented with fetal bovine serum, penicillin/streptomycin solution, and astrocyte growth supplement according to the manufacturer’s instructions, and cryopreserved as passage 2 working stocks. For experiments, astrocytes were thawed from passage 2 working stocks and directly seeded onto 96-well plates without further passaging.

All experiments were performed using 96-well black/clear-bottom poly-D-lysine-coated plates (Thermo Scientific, Cat. No. 152037). Plates were further coated with laminin 521 (BioLamina, Cat. No. BLA-LN521-05) at 0.5 µg/well overnight at 4°C under light-protected conditions. After removal of the laminin solution, astrocytes were seeded at 6,500 cells/well in 200 µL/well of complete astrocyte medium and maintained in a humidified incubator at 37°C with 5% CO₂ for at least 16 h. To reduce serum-containing medium before stimulation, the culture volume was adjusted to 280 µL/well and subjected to three sequential 4-fold medium exchanges. In each exchange, 210 µL of medium was removed and replaced with 210 µL of serum-free Astrocyte Medium. This procedure resulted in an approximately 64-fold dilution of the original serum-containing medium. Cells were then maintained in serum-free Astrocyte Medium for an additional 24 h before stimulation. To induce an inflammatory reactive astrocyte condition, cells were treated for 24 h with an A1-inducing cytokine/complement cocktail consisting of recombinant human IL-1α (Proteintech, Cat. No. HZ-1320), recombinant human TNF-α (Proteintech, Cat. No. HZ-1014), and human C1q (Hycult Biotech, Cat. No. HC2123-1MG) at final concentrations of 3 ng/mL, 30 ng/mL, and 400 ng/mL, respectively. For cytokine/complement withdrawal experiments, astrocytes were first exposed to the A1-inducing cocktail for 24 h. The stimulus-containing medium was then diluted by five sequential 4-fold medium exchanges using serum-free Astrocyte Medium. In each washout step, 210 µL of medium was removed from a total volume of 280 µL/well and replaced with 210 µL of fresh serum-free Astrocyte Medium. This procedure resulted in an approximately 1,024-fold dilution of the A1-inducing cytokine/complement cocktail. Cells were subsequently maintained in serum-free Astrocyte Medium for the indicated post-washout periods. For mock-control experiments, the same serial dilution procedure was performed in wells without astrocytes, and the resulting mock-treated medium was added to untreated astrocytes. For JAK inhibition experiments, baricitinib phosphate (Selleck, Cat. No. S5754) was prepared as a 20× stock solution and added to the culture medium at the indicated concentrations and time points. Vehicle-treated cultures were used as controls where applicable.

### RNA sequencing and TPM-based expression analysis

Total RNA was extracted from cultured human astrocytes using TRIzol reagent (Invitrogen) and the Direct-zol RNA Microprep kit (Zymo Research) according to the manufacturer’s instructions. Briefly, astrocytes were lysed directly in TRIzol after removal of culture medium. During purification, on-column DNase digestion was performed to remove genomic DNA. RNA was eluted in nuclease-free water and stored at −80°C. RNA concentration was assessed using a NanoDrop, and RNA integrity was evaluated using an Agilent TapeStation system with RNA ScreenTape. Samples with sufficient RNA quality were submitted to Azenta for RNA sequencing. Libraries were sequenced using the NovaSeq platform.

FASTQ files were evaluated using FastQC and processed using Trimmomatic version 0.36 to remove adaptor sequences and low-quality bases. Filtered reads were mapped to the human reference genome GRCh38 with Ensembl release 88 transcript annotation using STAR version 2.5.2b. Gene expression levels were estimated as transcripts per million (TPM) using RSEM version 1.2.31. For gene-level visualization, TPM values were used unless otherwise indicated. For selected genes, expression values were plotted as TPM or log₂(TPM + 1) values, as indicated in the corresponding figure legends.

### Principal component analysis

Principal component analysis (PCA) was performed using Python. Gene-level TPM values were first processed by collapsing duplicated gene symbols using the mean TPM value. Genes with TPM > 1 in at least two samples were retained for PCA. The filtered TPM matrix was transformed as log₂(TPM + 1), and PCA was performed using scikit-learn.

### Immunostaining

Cells were cultured as described in the cell culture section. Cells were fixed with fresh 4% paraformaldehyde in PBS and washed three times with PBS. After blocking with BSA-PBST containing 1% BSA and 0.1% Tween-20 for 10 min at room temperature, cells were incubated with anti-GFAP antibody (clone 2.2B10; Invitrogen; 1:400) in BSA-PBST for 1 h at room temperature. After washing three times with PBS, cells were incubated with Cy3-conjugated AffiniPure donkey anti-rat IgG (712-166-153; Jackson ImmunoResearch; 1:400) in BSA-PBST together with 5 µg/mL Hoechst 33258 for 1 h at room temperature in the dark. After staining, cells were washed three times with PBS, and images were acquired using a BIOREVO fluorescence microscope system (Keyence) and processed using ImageJ.

### Direct RT-qPCR analysis

At the indicated time points, conditioned media were collected and stored for downstream protein analysis. Astrocytes were then lysed directly in culture wells, and RT-qPCR was performed using the CellAmp™ Direct TB Green® RT-qPCR Kit (Takara Bio, Cat. No. 3737A) according to the manufacturer’s protocol. Quantitative PCR was performed using a QuantStudio 3 Real-Time PCR System (Applied Biosystems). Primer sequences used for quantitative PCR analysis are listed in Supplementary Table 1.

Gene expression levels were normalized to glyceraldehyde-3-phosphate dehydrogenase (GAPDH). ΔCt values were calculated as Ct_target − Ct_GAPDH, and normalized expression values were shown as −ΔCt where indicated. For relative expression analysis, fold changes were calculated using the 2^−ΔΔCt method. Samples with Ct values of 35 or greater, or with no detectable amplification, were considered non-detected and are indicated as N.D. in the corresponding figures.

### CD44 aptamer staining and fluorescence imaging

Human primary astrocytes were cultured under untreated or A1-stimulated conditions as described above. Aptamer staining for cluster of differentiation 44 (CD44) was performed using a 5′-Alexa Fluor 647-labeled truncated CD44-Apt1-derived DNA aptamer, designated Apt-CD44, synthesized by Integrated DNA Technologies (IDT). Apt-CD44 was designed based on the reported CD44-Apt1 sequence and was used to minimize sequence-length differences from the inactive control aptamer. The sequence of aptamers used in this study are listed in Supplementary Table 2. Before cell staining, the 20× aptamer solution was prepared in 1× PBSM (PBS containing 1 mM MgCl₂), denatured at 90°C for 3 min, and gradually cooled to 20°C to allow aptamer refolding. Five microliters of the folded 20× aptamer solution was added to 95 µL of astrocyte culture medium in each well without medium replacement. Cells were incubated with Apt-CD44 at a working concentration of 50 nM for 24 h at 37°C in a humidified incubator with 5% CO₂. After 24 h of aptamer incubation, Hoechst 33258 solution (Dojindo) was added directly to the culture medium at a final concentration of 1 µg/mL. Cells were further incubated for 1 h to label nuclei.

To remove unbound and nonspecifically associated aptamer, cells were washed by three sequential four-fold dilution washes with serum-free medium containing 0.1% dextran sulfate, followed by three additional four-fold dilution washes with PBS. Fluorescence images were acquired after washing using a BIOREVO fluorescence microscope system (Keyence). Alexa Fluor 647 and Hoechst signals were imaged using identical acquisition settings across experimental conditions. Representative images were obtained from multiple fields per well. Image visualization was performed using ImageJ/Fiji when applicable. Fluorescence microscopy experiments were performed in three independent experiments.

### Flow cytometric analysis of CD44 aptamer staining

For quantitative analysis of CD44 aptamer staining, human primary astrocytes were stained with Alexa Fluor 647-labeled Apt-CD44 at a working concentration of 100 nM using the same staining and washing procedures described in the CD44 aptamer staining and fluorescence imaging section. After washing, cells were detached using Accutase by incubation at 37°C for 5 min. Detached cells were gently dissociated by pipetting, collected, and kept on ice until flow cytometric analysis. To obtain sufficient cell numbers, cells from three wells under the same condition were pooled as one sample.

Alexa Fluor 647 fluorescence was analyzed using a BD LSRFortessa flow cytometer. Flow cytometry data were processed using FlowJo software. Cells were gated based on forward scatter and side scatter to exclude debris, followed by doublet exclusion to select single cells. Aptamer staining intensity was quantified as the mean fluorescence intensity (MFI) of the Alexa Fluor 647 signal in the single-cell population. Flow cytometry experiments were performed in three independent experiments, with each sample prepared by pooling three wells per condition.

### Western blotting analysis

Cell lysates were prepared using M-PER Mammalian Protein Extraction Reagent (Thermo Scientific), and protein concentrations were determined before SDS-PAGE analysis. Lysate samples were mixed with SDS sample buffer containing DTT and heated at 95°C for 3 min. For cellular protein analysis, 0.5 µg and 0.3 µg of total protein per lane were loaded for C3 and CD44 detection, respectively. Conditioned-media samples were mixed directly with SDS sample buffer, and equal volumes of medium were loaded unless otherwise indicated.

Samples were separated by SDS-PAGE using Mini-PROTEAN TGX Stain-Free gels. Stain-free gel images were acquired using a ChemiDoc Touch MP imaging system (Bio-Rad) before transfer. Proteins were transferred onto low-fluorescence PVDF membranes using a Trans-Blot Turbo Transfer System. Membranes were washed with PBST, blocked with PVDF Blocking Reagent for Can Get Signal (TOYOBO) for at least 60 min at room temperature, and incubated with primary antibodies against C3 (ab200999; Abcam; 1:1000), CD44 (ab157107; Abcam; 1:1000), GFAP (13-0300; Invitrogen; 1:1000), or GAPDH (60004-1-Ig; Proteintech; 1:1000) diluted in Can Get Signal Immunoreaction Solution 1 for 90 min at room temperature. After washing with PBST, membranes were then incubated with fluorescently labeled secondary antibodies diluted in Can Get Signal Immunoreaction Solution 2 for 60 min at room temperature. Mouse primary antibodies were detected using StarBright Blue 700-conjugated goat anti-mouse IgG (12004158; Bio-Rad; 1:2000), rabbit primary antibodies using StarBright Blue 520-conjugated goat anti-rabbit IgG (12005869; Bio-Rad; 1:2000), and rat primary antibodies using Cy3-conjugated AffiniPure donkey anti-rat IgG (712-166-153; Jackson ImmunoResearch; 1:800). Fluorescent signals were detected using a ChemiDoc Touch MP imaging system. Western blot images shown in the main figures were cropped to the relevant molecular weight ranges. The corresponding whole blot images are provided in Supplementary Information.

## RESULTS

### A1 stimulation induces an inflammatory transcriptional response and C3 protein expression in human primary astrocytes

To establish an in vitro model of inflammatory reactive astrocyte activation, human primary astrocytes were left untreated (UT) or exposed to A1 stimulation, defined here as treatment with TNF-α, IL-1α, and C1q for 24 h under serum-free culture conditions (Fig. 1a). To assess astrocyte morphology and overall cell integrity under this experimental setting, immunocytochemical analysis was performed. Immunostaining for glial fibrillary acidic protein (GFAP) showed that astrocyte-like morphology was largely preserved in both UT and A1-stimulated cultures, with no obvious cell loss or morphological deterioration after A1 stimulation (Fig. 1b).

**Figure 1.**
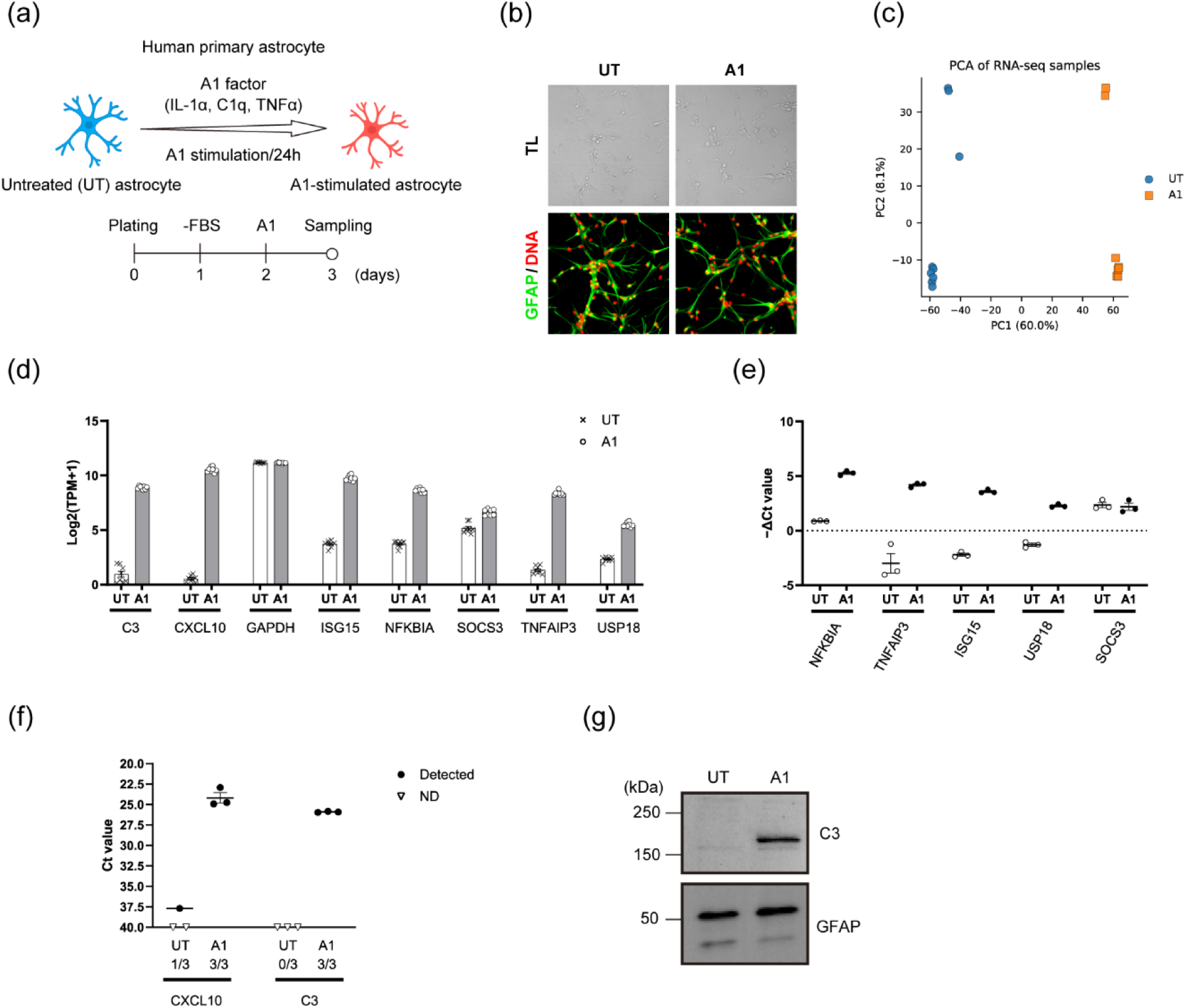
A1 stimulation induces inflammatory transcriptional responses and C3 protein expression in human primary astrocytes. **(a)** Schematic overview of the experimental design. Human primary astrocytes were cultured under serum-free conditions and stimulated with A1 stimulation consisting of IL-1α, C1q, and TNFα for 24 h. **(b)** Representative transmitted-light and immunofluorescence images of untreated and A1-stimulated astrocytes. Cells were stained for GFAP and DNA. **(c)** Principal component analysis of RNA-seq samples from untreated and A1-stimulated astrocytes. PCA was performed using log₂(TPM + 1)-transformed expression values after filtering expressed genes. **(d)** RNA-seq analysis showing expression levels of selected inflammatory and astrocyte-associated genes in untreated and A1-stimulated astrocytes. Expression values are shown as log₂(TPM + 1). **(e)** qPCR validation of selected inflammatory response genes following A1 stimulation. Expression values were normalized to GAPDH and are shown as −ΔCt values. **(f)** Raw Ct values and detection status for CXCL10 and C3 in untreated and A1-stimulated astrocytes. Non-detected samples are indicated as ND. The fractions below each condition indicate the number of detected samples over the total number of replicates (detected/total). **(g)** Representative western blot showing detection of C3 protein in A1-stimulated astrocytes. GFAP was used as an astrocyte-associated protein control. Data points represent biological replicate samples; bars indicate mean ± SEM where applicable. RNA-seq data were obtained from n = 8 independent well-derived samples per condition. qPCR data were obtained from n = 3 independent well-derived samples per condition. Data points in qPCR panels represent independent well-derived samples; bars indicate mean ± SEM where applicable. Immunofluorescence and western blot images are representative of at least 3 independent experiments. ND indicates not detected. Uncropped western blot images are provided in Supplementary Fig. 5.

We first examined whether A1 stimulation altered the global transcriptional state of cultured human astrocytes. Principal component analysis of RNA-seq samples showed a clear separation between untreated and A1-stimulated astrocytes along PC1, indicating that A1 stimulation induced a robust transcriptional remodeling of human primary astrocytes (Fig. 1c). Consistent with this global shift, RNA-seq analysis showed increased expression of inflammatory and reactive astrocyte-associated genes, including C3, CXCL10, ISG15, NFKBIA, TNFAIP3, and USP18, following A1 stimulation (Fig. 1d). Of note, these RNA-seq data were obtained from two independent human astrocyte lots. PCA showed that samples separated primarily by treatment condition rather than by astrocyte lot, and similar induction patterns of inflammatory and reactive astrocyte-associated genes were observed across the two lots.

We next validated selected inflammatory response genes by qPCR. A1 stimulation markedly increased the expression of NF-κB-associated genes, including NFKBIA and TNFAIP3, as well as interferon/inflammatory response genes such as ISG15 and USP18 (Fig. 1e). CXCL10 and C3 were undetectable or near the detection limit in untreated astrocytes but became clearly detectable after A1 stimulation, as shown by raw Ct values and detection frequency (Fig. 1f). Similar induction of selected inflammatory response genes was observed in an independent astrocyte lot, supporting the reproducibility of the A1-stimulated response across astrocyte preparations (Supplementary Fig. 1). At the protein level, western blot analysis showed that C3 protein bands were not detected or were barely detectable in UT astrocytes but were clearly induced after A1 stimulation, consistent with the qPCR results (Fig. 1g).

Together, these results confirm that A1 stimulation induces a robust inflammatory transcriptional response in cultured human primary astrocytes and establish C3 as a strongly inducible complement-associated readout in this system.

### CD44-based analysis provides an orthogonal phenotypic readout of A1-stimulated astrocyte remodeling

Because cultured human astrocytes exhibited basal GFAP expression under UT conditions, GFAP alone may not fully capture A1-stimulated phenotypic remodeling. We therefore examined whether A1 stimulation altered astrocyte phenotype at the cell-surface protein level by assessing CD44 protein expression and CD44-targeting aptamer binding as orthogonal phenotypic readouts.

CD44 is a hyaluronan-binding cell-surface glycoprotein originally identified as a principal hyaluronan receptor and has been implicated in astrocyte phenotypes, including reactive and disease-associated astrocyte states [19–21]. RNA-seq analysis showed that CD44 was expressed in UT human primary astrocytes and was modestly increased after A1 stimulation (Fig. 2a). Consistent with this basal mRNA expression, western blotting with a pan-CD44 antibody detected a major CD44-immunoreactive band at approximately 75–80 kDa in both UT and A1-stimulated astrocytes (Fig. 2b). This band size is compatible with the expected migration range of the standard CD44 isoform, CD44s, which lacks variant exons (Fig. 2b) [19, 22]. In addition, A1 stimulation increased a broad high-molecular-weight CD44-immunoreactive smear around 200 kDa, hereafter referred to as CD44-HMW (Fig. 2b). CD44s protein levels were only modestly changed, whereas the CD44-HMW signal was more clearly enhanced after A1 stimulation (Fig. 2b, c). In contrast, GFAP protein levels were not significantly altered under the same conditions (Fig. 2b, c). These findings suggest that A1 stimulation does not simply increase total CD44 abundance, but is associated with altered CD44-related protein band patterns, indicating surface-associated phenotypic remodeling.

**Figure 2.**
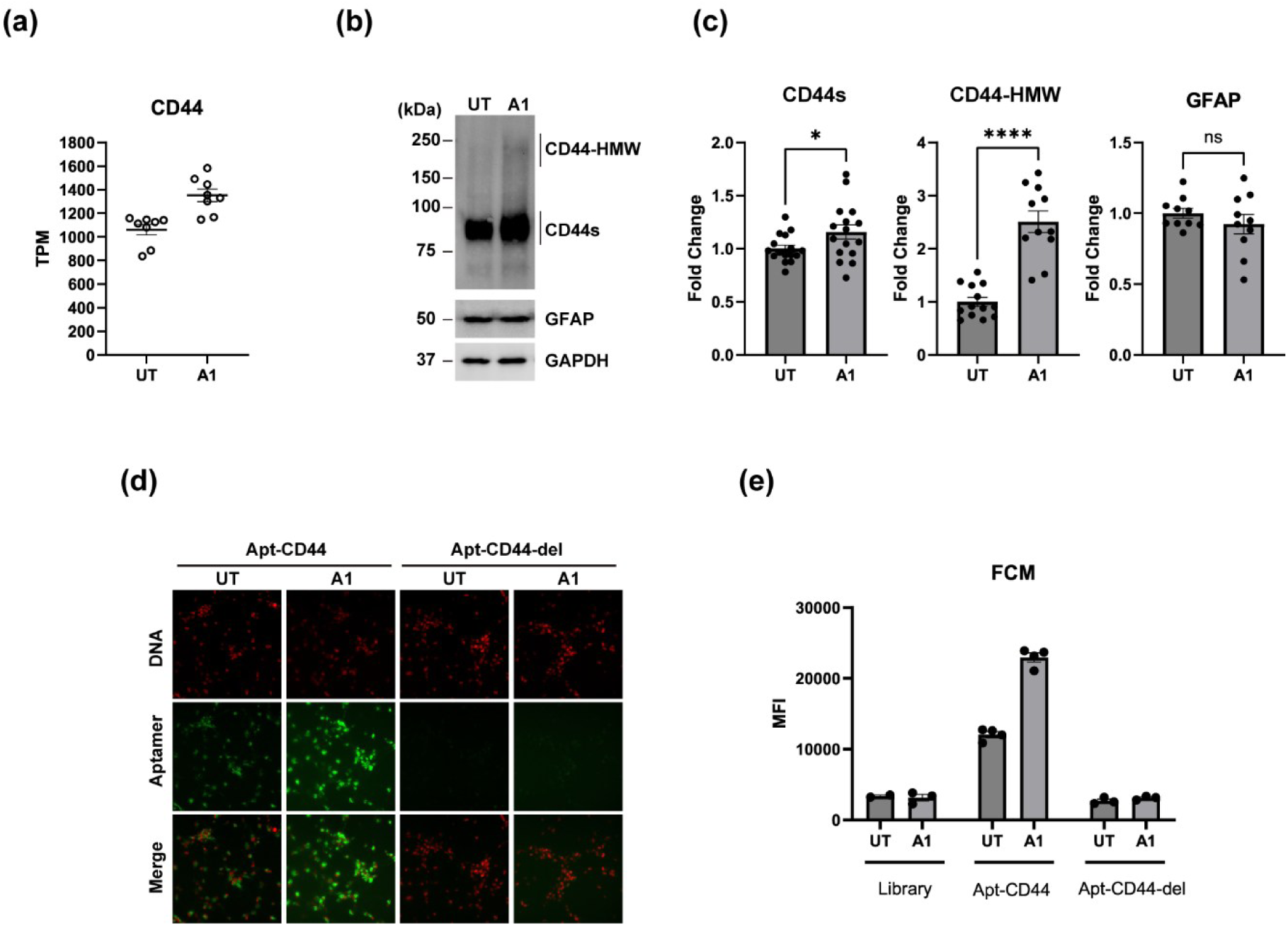
CD44 aptamer binding provides an orthogonal phenotypic readout of A1-induced astrocyte remodeling. **(a)** RNA-seq expression of CD44 in untreated and A1-stimulated human primary astrocytes. Expression values are shown as TPM. **(b)** Representative western blot analysis of CD44 isoform-associated protein bands, GFAP, and GAPDH in untreated and A1-stimulated astrocytes. CD44s and CD44-HMW indicate standard CD44 and high-molecular-weight CD44 bands, respectively. **(c)** Quantification of CD44s, CD44-HMW, and GFAP protein levels from western blot analyses. Values are shown as fold change relative to untreated controls. **(d)** Representative fluorescence images showing binding/internalization of Apt-CD44 and the inactive deletion control aptamer, Apt-CD44-del, to untreated and A1-stimulated astrocytes. DNA staining is shown as a nuclear counterstain. **(e)** Flow cytometric analysis of aptamer binding/internalization in untreated and A1-stimulated astrocytes. Mean fluorescence intensity (MFI) is shown for the aptamer library, Apt-CD44, and Apt-CD44-del. RNA-seq data are from the dataset described in Figure 1. Western blot quantification was based on independent well-derived samples from 4 independent experiments. Representative western blot and aptamer staining images are shown from at least 3 independent experiments. Flow cytometry was performed in n = 3 independent experiments. Bars indicate mean ± SEM; statistical comparisons are shown where indicated. Uncropped western blot images are provided in Supplementary Fig. 6.

We next evaluated whether this altered cell surface phenotype could be detected using a CD44-binding DNA aptamer, CD44-Apt1 [23]. This aptamer was originally identified as a CD44E/CD44s dual-targeting aptamer and was reported to bind both CD44E and CD44s with high affinity and specificity, while showing little association with the hyaluronan-binding domain of CD44 [23]. We therefore used a 5′-Alexa Fluor 647-labeled truncated version of CD44-Apt1-derived aptamers (hereafter, Apt-CD44, Supplementary Table 2). The binding activity of Apt-CD44 series was evaluated by performing aptamer pull-down followed by western blotting using 5′-biotinylated aptamers. Pull-down assays confirmed that Apt-CD44 recovered CD44-immunoreactive proteins from HepG2 and astrocyte lysates, whereas Apt-CD44-del showed reduced recovery (Supplementary Fig. 2). Notably, high-molecular-weight CD44 species were preferentially recovered from A1-stimulated astrocyte lysates, supporting the possibility that A1 stimulation alters CD44-associated molecular states recognized by Apt-CD44 (Supplementary Fig. 2).

for fluorescence imaging and flow cytometric analysis of human primary astrocytes. Fluorescence imaging showed increased Apt-CD44 signal in A1-stimulated astrocytes compared with UT astrocytes, whereas an inactive deletion control aptamer, Apt-CD44-del, showed minimal binding under both conditions (Fig. 2d). Imaging analysis revealed enhanced cellular internalization activity of Apt-CD44 in A1-stimulated astrocytes (Fig. 2d). Flow cytometric analysis further confirmed increased Apt-CD44 binding/internalization activity after A1 stimulation, while the aptamer library control and inactivated Apt-CD44-del showed substantially lower signals (Fig. 2e and Supplementary Fig. 3).

Pull-down assays confirmed that Apt-CD44 recovered CD44-immunoreactive protein bands from HepG2 and astrocyte lysates. Notably, high-molecular-weight CD44 species were preferentially recovered from A1-stimulated astrocyte lysates, supporting the possibility that A1 stimulation alters CD44-associated molecular states recognized by Apt-CD44.

Together, these results indicate that A1 stimulation induces CD44-associated surface phenotypic alteration, characterized by increased high-molecular-weight CD44 species and enhanced CD44 aptamer internalization activity, while GFAP expression remains largely unchanged. Thus, CD44 aptamer analysis provides an orthogonal phenotypic assay supporting the interpretation that A1 stimulation alters astrocyte state beyond transcriptional inflammatory gene responses.

### C3 expression persists after withdrawal of inflammatory stimulation

We next investigated whether the inflammatory astrocyte state was reversible after removal of the inflammatory stimulus. Human primary astrocytes were treated with A1 stimulation and then subjected to stimulus washout, followed by time-course analysis of inflammatory gene expression (Fig. 3a). Time labels indicate the duration of each indicated culture condition; for washout samples, the indicated time represents the duration after removal of A1 stimulation.

**Figure 3.**
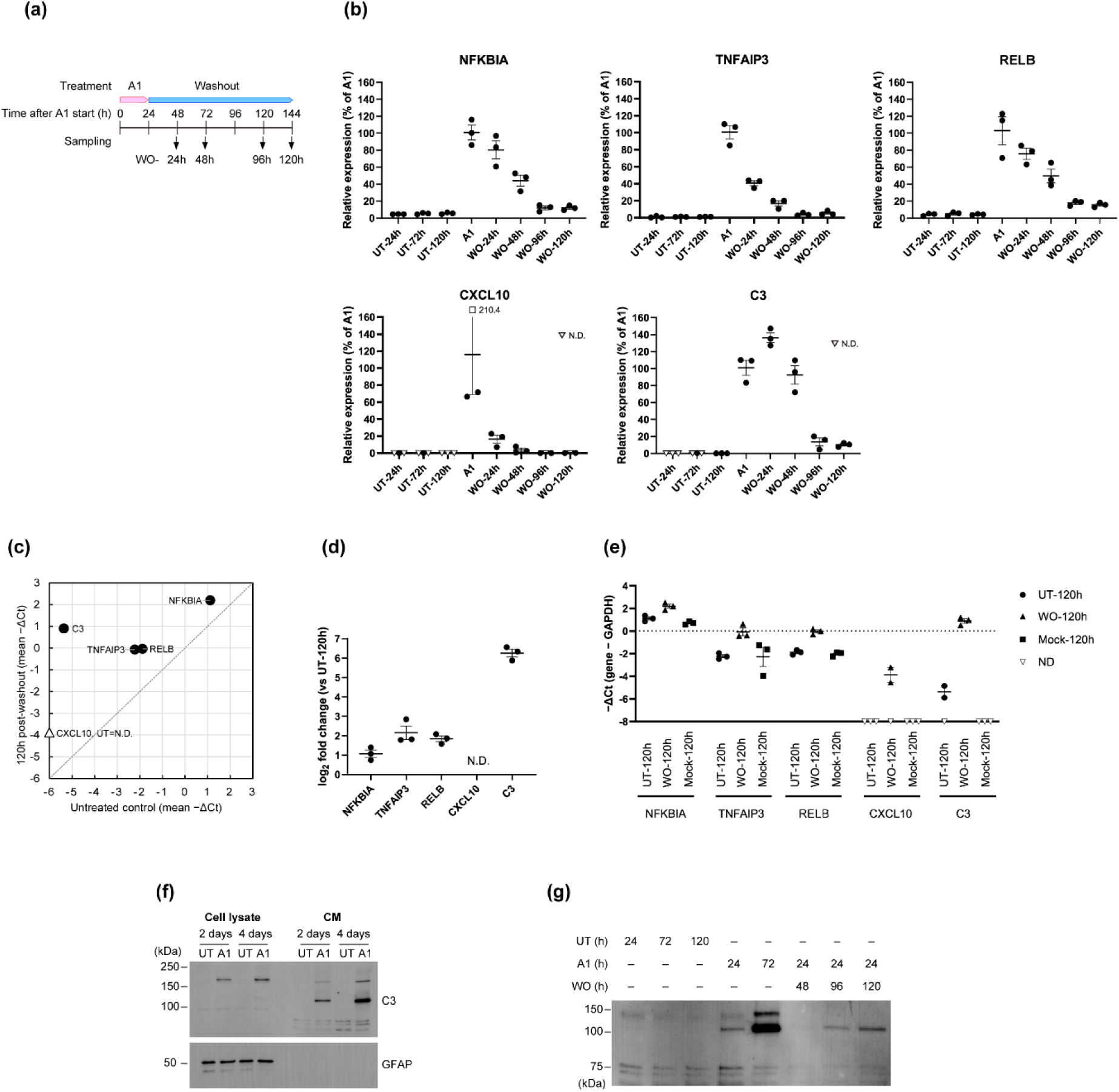
C3 expression persists after withdrawal of A1 stimulation. **(a)** Schematic overview of the A1 stimulation and washout time-course experiment. Human primary astrocytes were either maintained under untreated conditions (UT), continuously exposed to A1 stimulation (A1), or exposed to A1 stimulation followed by cytokine washout (WO). Time labels indicate the duration of each indicated culture condition. For washout samples, time indicates the duration after removal of A1 stimulation. **(b)** Time-course analysis of inflammatory gene expression after A1 stimulation and cytokine washout. Expression values were normalized to GAPDH and are shown as percentage of the A1-24h condition for each gene. Non-detected samples are indicated as N.D. **(c)** Scatter plot comparing mean GAPDH-normalized expression values in untreated control and 120 h post-washout samples. Values are shown as mean −ΔCt. The diagonal line indicates equal expression between untreated and post-washout conditions. CXCL10 was non-detected in untreated samples and is indicated accordingly. **(d)** Fold change in gene expression at 120 h post-washout relative to time-matched untreated controls. Fold changes were calculated from GAPDH-normalized qPCR values. C3 expression in untreated controls was near the assay detection limit; therefore, fold-change values were interpreted together with −ΔCt values and detection status. **(e)** GAPDH-normalized expression of selected genes in untreated, mock-treated, and 120 h post-washout astrocytes. Values are shown as −ΔCt. Non-detected samples are indicated as N.D. **(f)** Representative western blot analysis of C3 protein bands in cell lysates and conditioned medium from untreated and A1-stimulated astrocytes at the indicated time points. Equal sample volumes were loaded for conditioned medium. **(g)** Representative western blot analysis of conditioned medium showing detection of C3 protein bands after A1 stimulation and after cytokine washout. Condition labels indicate the duration of each indicated culture condition; for washout samples, time indicates the duration after removal of A1 stimulation. The conditioned-medium blot is shown as a qualitative detection assay rather than a quantitative comparison of secreted C3 levels. Data points represent biological replicate samples; bars indicate mean ± SEM. qPCR data were obtained from n = 3 independent well-derived samples per condition. Western blot images are representative of at least 3 independent experiments. Data points represent independent well-derived samples; bars indicate mean ± SEM where applicable. ND indicates not detected. Uncropped western blot images are provided in Supplementary Fig. 7.

When expression values were normalized to the 24 hours of A1 stimulation (A1), NF-κB-associated genes such as NFKBIA, TNFAIP3, and RELB were strongly induced by A1 stimulation but progressively decreased after stimulus washout (Fig. 3b). CXCL10 was also induced by A1 stimulation and rapidly declined by 24 h after washout (Fig. 3b). In contrast, C3 showed a distinct pattern. C3 expression remained high at 24 h and 48 h after stimulus washout, and detectable expression was maintained at 96 h and later post-washout time points (Fig. 3b).

To compare post-washout expression with time-matched untreated controls, we plotted mean −ΔCt values in untreated and 120 h post-washout astrocytes. Most NF-κB-associated genes were positioned close to the diagonal line, indicating that their expression after washout returned toward untreated levels (Fig. 3c). In contrast, C3 deviated from this pattern and remained elevated in post-washout astrocytes relative to untreated controls (Fig. 3c). Consistently, fold-change analysis relative to time-matched untreated controls showed that C3 remained strongly elevated at 120 h post-washout, whereas other inflammatory response genes showed smaller changes (Fig. 3d). Because C3 expression in untreated controls was near the assay detection limit, fold-change values were interpreted together with GAPDH-normalized −ΔCt values and detection status to avoid overinterpretation of changes caused by low baseline expression. Mock-treated controls showed no detectable effect of residual A1 factors after washout. These results further supported the conclusion that C3 remained detectably elevated after stimulus withdrawal, whereas CXCL10 was largely undetected and NF-κB-associated genes were closer to untreated or mock-treated levels (Fig. 3e). A similar pattern of sustained C3 expression after A1 stimulus withdrawal was also observed in an independent astrocyte lot, indicating that this response was not restricted to a single astrocyte preparation (Supplementary Fig. 4).

We next analyzed cellular and conditioned-medium C3 protein levels by western blotting. In cell lysates, C3-related protein bands were detected after 2 and 4 days of A1 stimulation, consistent with the induction of C3 mRNA (Fig. 3f). In conditioned medium (CM), C3-related protein bands were also detected after 2 and 4 days of A1 stimulation, indicating that C3 protein was released into the culture medium (Fig. 3f). The major cellular C3-related band was detected at approximately 180 kDa, whereas the conditioned-medium C3-related band was detected at approximately 110 kDa, suggesting that C3 was present in a processed or cleaved form in the culture medium (Fig. 3f).

We next examined whether transient A1 stimulation led to detectable C3-related proteins in conditioned medium after stimulus washout. C3-related protein bands were not detected in conditioned medium from untreated astrocyte cultures even after 120 h. In contrast, 24 h of transient A1 stimulation induced detectable C3 protein release. After stimulus washout, C3-related protein bands were not clearly detected at 48 h but became detectable in conditioned medium at 96 h and later time points, suggesting gradual accumulation of C3-related proteins in the culture medium (Fig. 3g). Because conditioned-medium samples were analyzed as a qualitative assay using equal sample volumes rather than a normalized quantitative assay, these data were interpreted as evidence for the presence of detectable C3-related protein in the culture medium after washout.

Together, these findings show that A1-induced inflammatory responses are not uniformly persistent after stimulus withdrawal. In particular, C3 showed distinct post-stimulus persistence compared with other transient inflammatory-response genes, supporting the presence of a post-inflammatory C3-high astrocyte state.

### JAK inhibition attenuates persistent C3 expression after A1 stimulus withdrawal

Because C3 gene expression remained elevated after A1 stimulus washout, we next asked whether this persistent C3-high state could be pharmacologically attenuated. We focused on JAK signaling because A1 stimulation induced multiple inflammatory and interferon-associated response genes, and because JAK/STAT signaling is known to contribute to reactive astrocyte regulation [17, 24].

To examine whether the persistent C3-high state could be attenuated by JAK inhibition, we treated post-washout astrocytes with baricitinib, a JAK1/2 inhibitor [25]. Astrocytes were first exposed to A1 stimulation, subjected to stimulus washout, and then treated with baricitinib during the post-washout phase. In 120 h post-washout astrocytes, persistent C3 expression was reduced by baricitinib in a concentration-dependent manner (Fig. 4a). In contrast, the effects of baricitinib on NFKBIA, TNFAIP3, and RELB were more modest or variable, consistent with the observation that these genes had already declined toward baseline after cytokine washout (Fig. 4a). SOCS3 expression was also reduced by baricitinib, supporting pharmacological suppression of JAK/STAT-associated signaling in this system (Fig. 4a). These results indicate that persistent C3 expression can be pharmacologically attenuated after removal of the initiating inflammatory stimulus.

**Figure 4.**
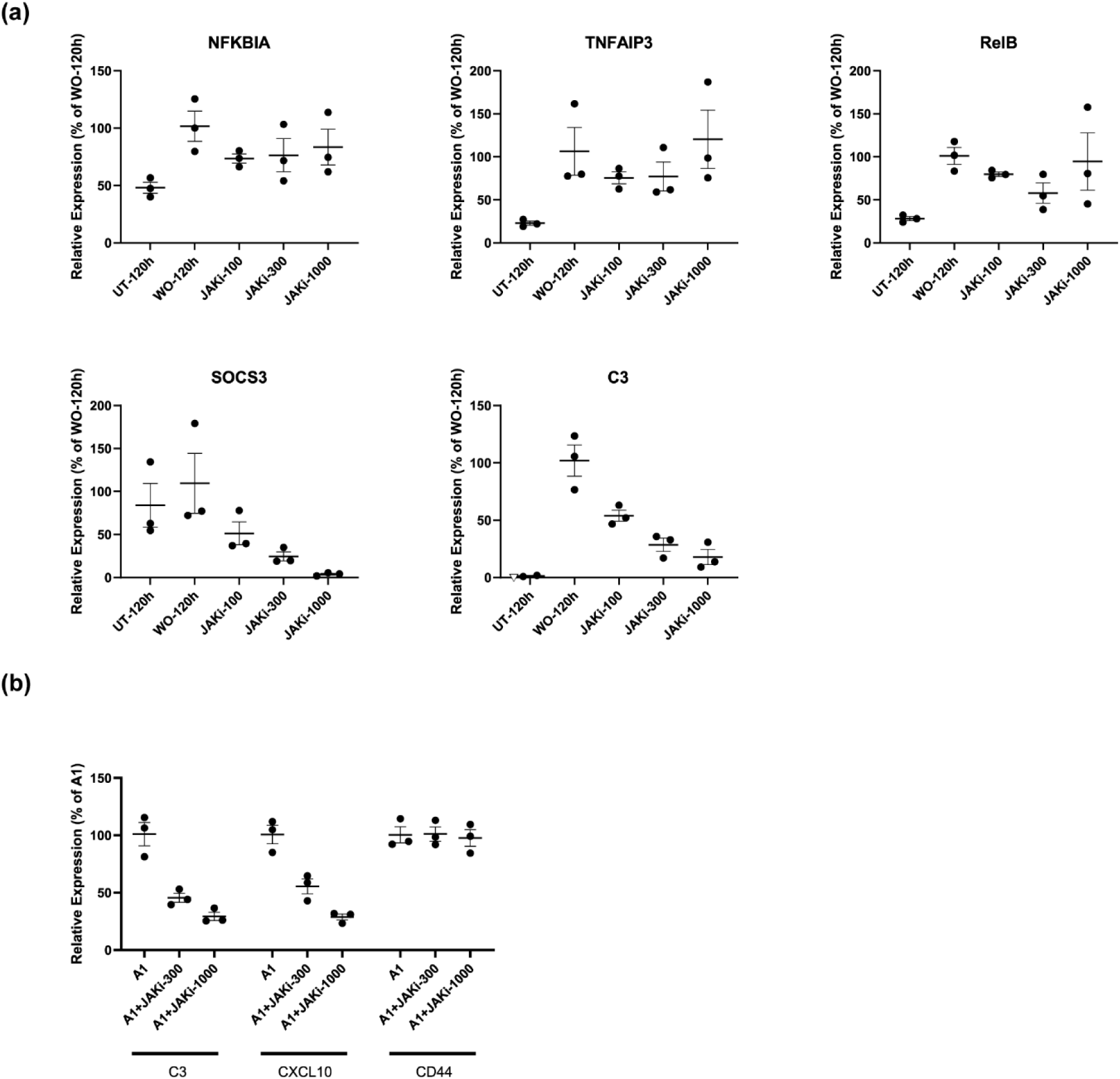
JAK inhibition attenuates persistent C3 expression after withdrawal of A1 stimulation. **(a)** qPCR analysis of reactive astrocyte-associated genes in time-matched untreated astrocytes (UT-120h), 120 h post-washout astrocytes (WO-120h), and post-washout astrocytes treated with 100 nM, 300 nM, and 1000 nM JAK inhibitor (JAKi-100, JAKi-300, JAKi-1000). Expression values were normalized to GAPDH and are shown as percentage of the mean WO-120h condition for each gene. JAK inhibitor was added after removal of A1 stimulation during the post-washout phase. **(b)** qPCR analysis of C3, CXCL10, and CD44 expression in A1-stimulated astrocytes (A1) and astrocytes stimulated with A1 in the presence of 300 nM and 1000 nM JAK inhibitor (A1+JAKi-300, A1+JAKi-1000). Expression values were normalized to GAPDH and are shown as percentage of the mean A1 condition for each gene. qPCR data were obtained from n = 3 independent well-derived samples per condition. Data points represent independent well-derived biological replicate samples; bars indicate mean ± SEM.

We also examined the effect of JAK inhibition on A1 stimulation. Baricitinib treatment reduced A1 stimulation-induced C3 and CXCL10 expression, whereas CD44 expression was less affected at the mRNA level (Fig. 4b). These results suggest that JAK signaling contributes to both inflammatory gene induction and the maintenance of persistent C3 expression after inflammatory stimulus withdrawal, although not all inflammatory phenotypic features appear to be equally JAK-dependent.

Together, these results support a model in which transient exposure to A1 stimulation generates a persistent C3-high astrocyte state that is maintained after broader inflammatory gene responses decline and can be attenuated by JAK inhibition.

## DISCUSSION

In this study, we found that transient exposure of astrocytes to A1 stimulation resulted in a residual inflammatory state in which C3 expression remained elevated after removal of the initial inflammatory stimulus, whereas other inflammatory-response genes, including CXCL10, ISG15, USP18, MX1, and NF-κB-related genes, returned toward baseline. These findings suggest that the post-stimulus astrocyte state is not simply a prolonged acute inflammatory response, but may represent a more selective residual reactive state. Importantly, persistent C3 expression was reduced by JAK inhibition, supporting a model in which JAK-dependent signaling contributes to the maintenance of this residual C3-high state after stimulus withdrawal.

C3 is not only a marker of inflammatory astrocyte activation, but has also been implicated as a functional complement mediator in astrocyte–neuron and astrocyte–microglia signaling, neuroinflammatory pathology, and synaptic dysfunction [26–28]. In particular, our findings are consistent with previous observations that astrocytic C3 can remain pathologically relevant beyond the initial inflammatory phase. In a recent laparotomy model, astrocytic C3 persisted after surgery even after several inflammatory cytokines had declined, and this persistent C3 was associated with sustained microglial activation, synaptic engulfment, and cognitive dysfunction [26]. Although our in vitro system does not directly test neuronal or synaptic functions, these observations support the possibility that persistence of astrocytic C3 after inflammatory stimulus withdrawal may represent a feature of post-inflammatory astrocyte responses rather than a simple experimental artifact.

More broadly, persistent C3 expression may have mechanistic links to microglia-mediated synapse elimination and neuroinflammatory pathology. Previous studies have shown that microglia can phagocytose complement-tagged synapses through C3-related complement pathways, and that astrocyte-derived C3 can facilitate microglial synaptic engulfment under inflammatory conditions [14, 29, 30]. These observations suggest that persistent astrocytic C3 may remain pathologically relevant even after other inflammatory factors have largely returned toward baseline levels.

Another important implication of this study is that JAK-STAT signaling may be associated not only with the induction of astrocyte reactivity, but also with maintenance of a post-inflammatory astrocyte state. JAK/STAT3 pathway has been known as a common inducer of astrocyte reactivity across disease models [17]. Suppression of astrocytic STAT3 signaling can reduce reactive astrogliosis and improve pathological and cognitive outcomes in Alzheimer’s disease models [31]. Our results support the possibility that JAK-dependent signaling may help to stabilize a persistent C3-high astrocyte state after removal of inflammatory stimulus. In this regard, JAK inhibition may be valuable not only for preventing inflammatory astrocyte induction, but also for suppressing persistent C3-driven astrocyte reactivity.

Our orthogonal analyses further support the interpretation that the A1-stimulated phenotype is not limited to acute inflammatory gene induction. In CD44-based analysis, A1 stimulation enhanced uptake of a CD44-targeting aptamer, indicating that this assay provided an additional surface-level phenotypic readout associated with the inflammatory astrocyte state. Thus, altered CD44 aptamer uptake in our model may reflect A1 stimulation-associated changes in astrocyte surface properties, complementing transcriptional inflammatory gene analyses.

This study has several limitations. First, our conclusions are based primarily on an in vitro astrocyte model, and additional in vivo validation will be important to determine how closely this persistent C3-high state reflects astrocyte behavior in disease-associated brain environments. Second, although JAK inhibition reduced persistent C3 expression, the precise upstream and downstream molecular mechanisms remain to be defined. Third, while astrocyte-derived or complement-associated C3 has been implicated in synaptic pathology in prior studies, we did not directly test neuronal or synaptic function in our system. Finally, it remains to be addressed whether the observed C3-high reactive state reflects persistent expression of selected genes or represents a distinct astrocyte state. Addressing these questions will be important in future work.

In summary, our findings support a model in which transient inflammatory stimulation induces a persistent C3-high residual reactive state in astrocytes, even after broader inflammatory genes have largely returned toward baseline. Because JAK inhibition suppressed this state, our findings suggest that its maintenance is pharmacologically modifiable. These results suggest that astrocyte inflammatory activity may persist beyond the initial inflammatory insult, potentially contributing to prolonged neuroinflammatory states. Therefore, targeting maintenance pathways may help limit persistent astrocyte reactivity, in addition to addressing the original inflammatory stimulus.

## LIST OF ABBREVIATIONS

C1q: Complement component 1q
C3: Complement component 3
CD44: Cluster of differentiation 44
CD44-HMW: High-molecular-weight
CD44: CD44s Standard CD44 isoform
CXCL10: C-X-C motif chemokine ligand 10
GAPDH: Glyceraldehyde-3-phosphate dehydrogenase
GFAP: Glial fibrillary acidic protein
IL-1α: Interleukin-1 alpha
JAK: Janus kinase
NF-κB: Nuclear factor kappa B
STAT: Signal transducer and activator of transcription
TNF-α: Tumor necrosis factor alpha
CM: Conditioned medium
CNS: Central nervous system
PCA: Principal component analysis
RSEM: RNA-Seq by Expectation-Maximization

## DECLARATIONS

### Author contributions

Y.S. conceived and led the study described in this manuscript, designed the experiments, analyzed the data, and drafted the manuscript. K. Okahara performed the experiments, including astrocyte culture, flow cytometry, and immunocytochemistry and analyzed data for the manuscript. J.K. supported astrocyte culture experiments. K.E. performed qPCR and western blotting experiments. Y.O. supported data analysis. K.Ohba contributed to project oversight and supported the study as the AMED project representative. All authors reviewed and approved the final manuscript.

### Availability of data and materials

The processed RNA-seq expression data supporting the findings reported in this article, including TPM values used for PCA and gene-level visualization, is available from the primary corresponding author (Y.S.) upon reasonable request and subject to institutional requirements. Other data supporting the findings of this study are available from the corresponding author upon reasonable request.

## Supplementary Information

**Supplementary Table 1.**
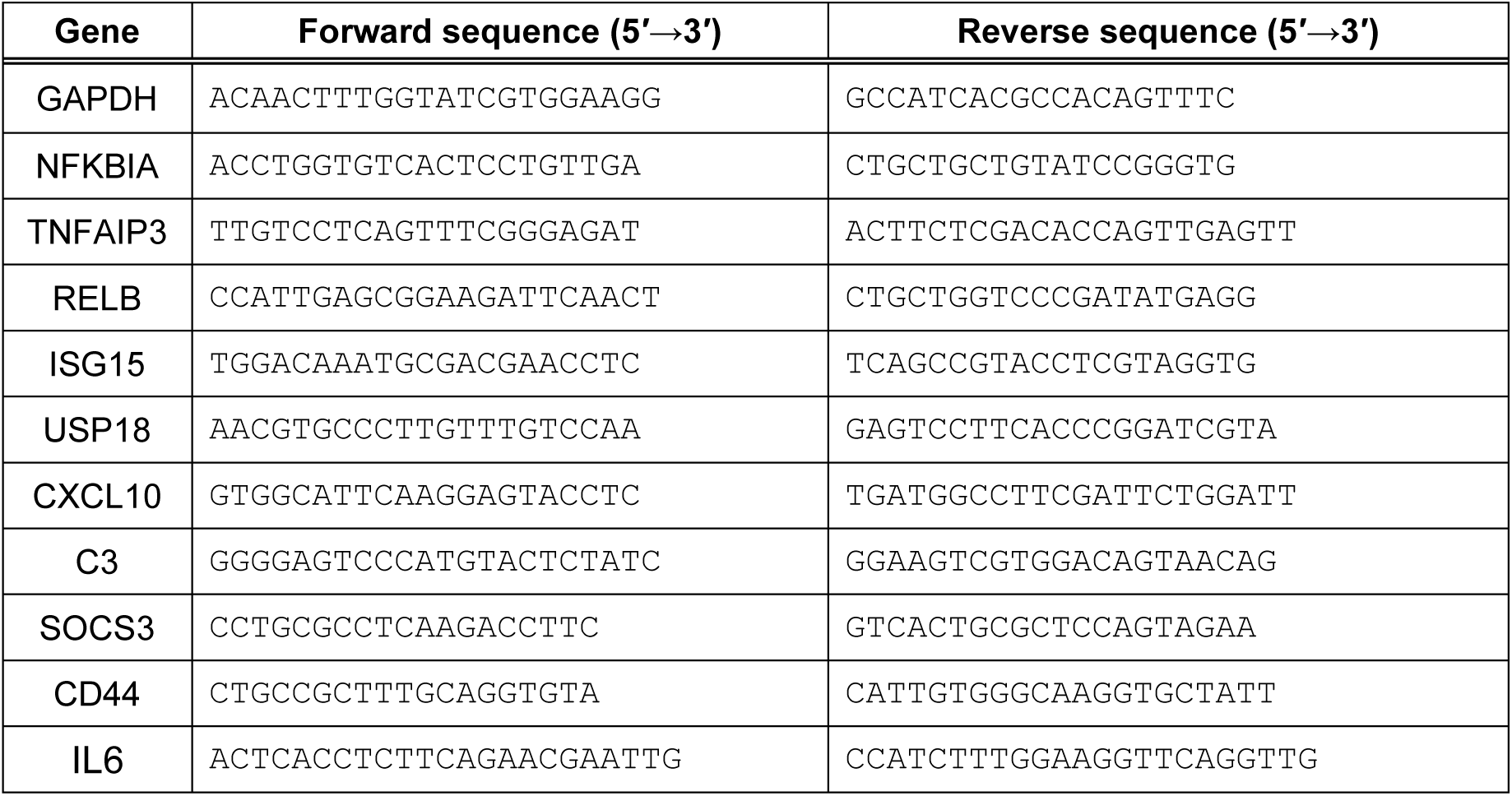
Primer sequences used in this study.

**Supplementary Table 2.**
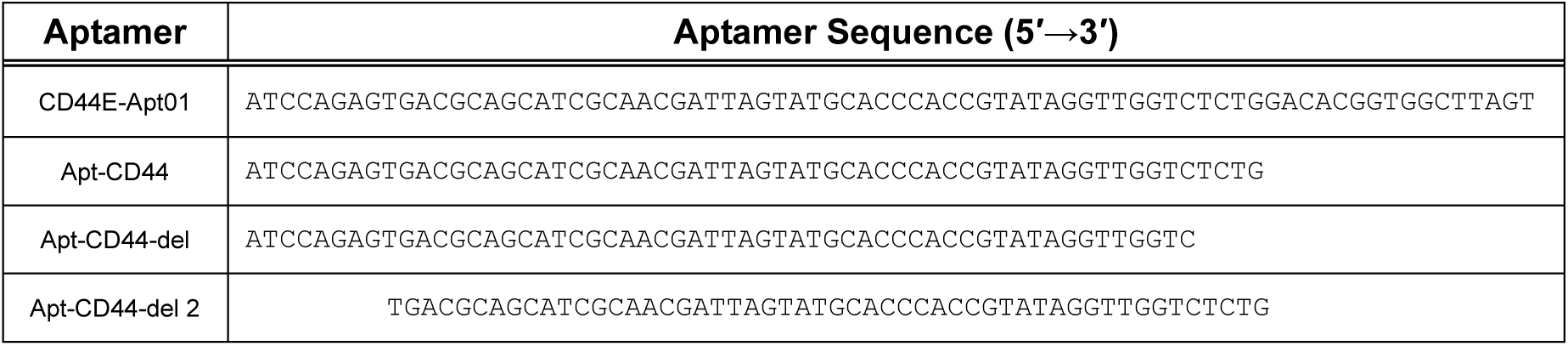
Sequences of CD44E-Apt01 and its truncated or deletion-derived DNA aptamers.

**Supplementary Figure 1.**
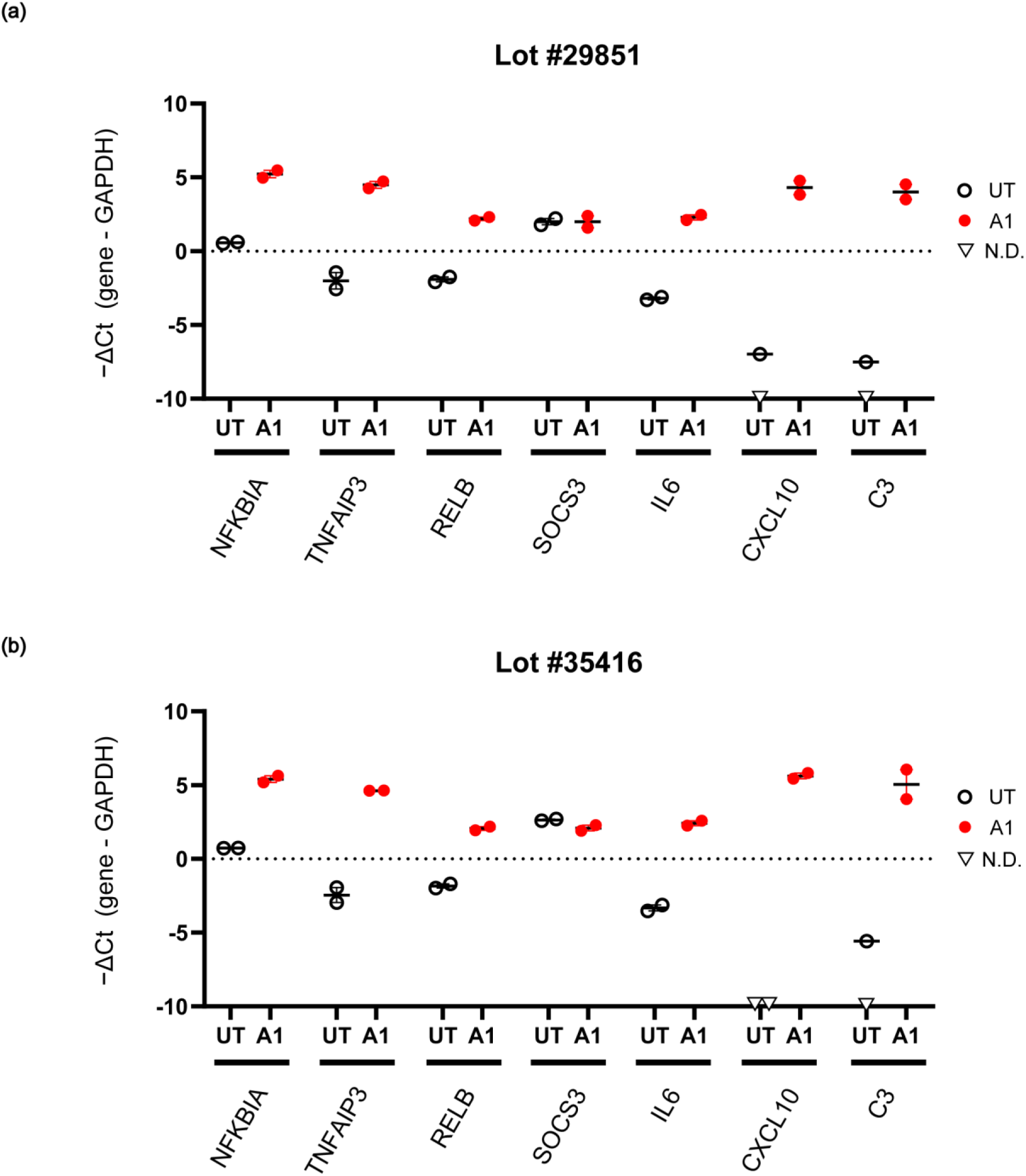
A1 stimulation induces similar inflammatory gene expression responses in independent astrocyte lots. qPCR validation of selected inflammatory response genes following A1 stimulation are shown for (a) lot 29851 and (b) lot 35416. Expression values were normalized to GAPDH and are shown as −ΔCt values.

**Supplementary Figure 2.**
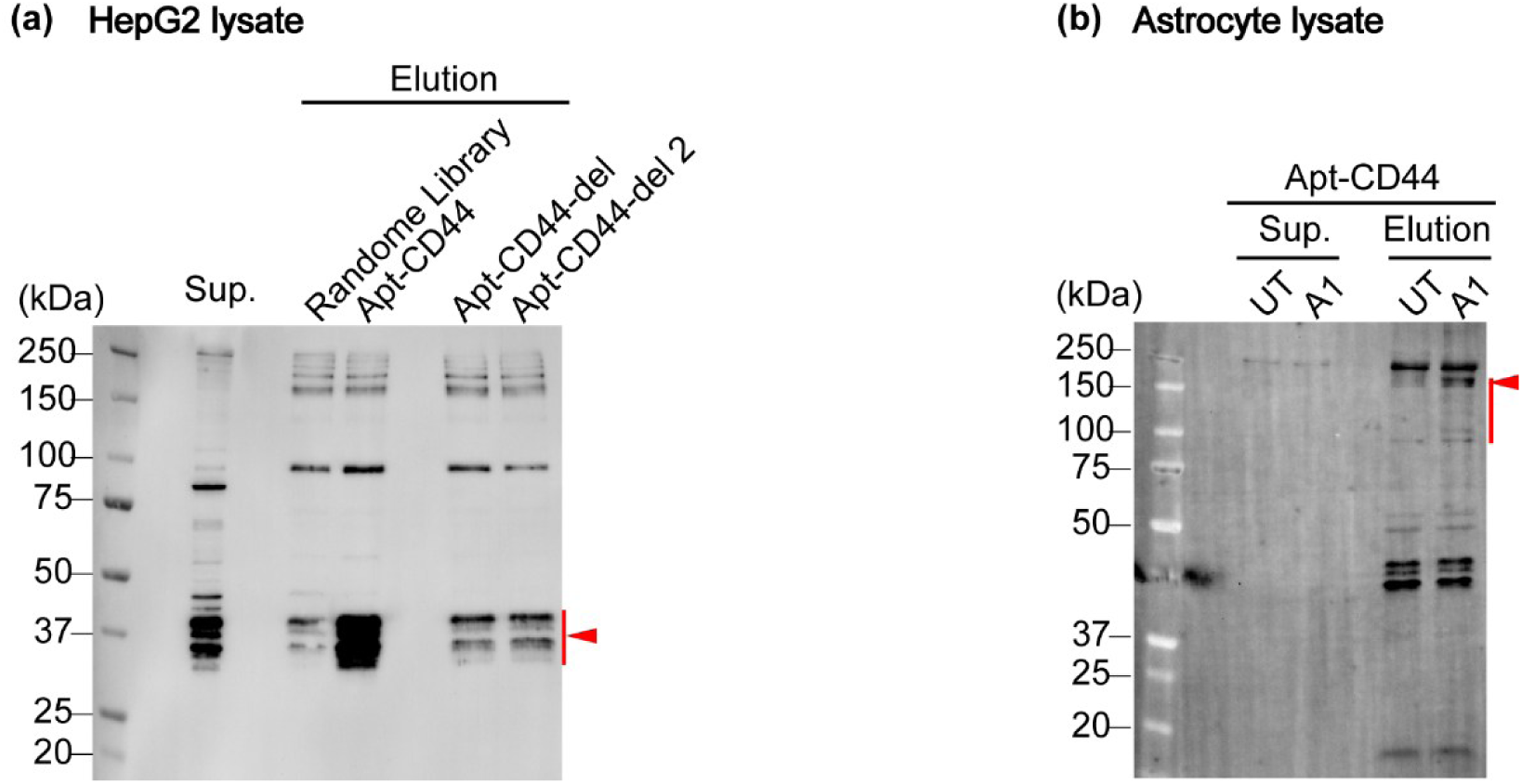
Apt-CD44 pull-down analysis of CD44-immunoreactive protein species. (a) CD44 aptamer pull-down assay using HepG2 cell lysate is shown for a control lysate. Eluted fractions were analyzed by western blotting for CD44. Apt-CD44 recovered CD44-immunoreactive bands, whereas the inactive control aptamers, Apt-CD44-del and Apt-CD44-del 2, showed reduced recovery of the lower-molecular-weight CD44 species indicated by the red arrowhead. (b) CD44 aptamer pull-down assay using untreated (UT) and A1-stimulated astrocyte lysates. Apt-CD44 recovered high-molecular-weight CD44 species from A1-stimulated astrocyte lysate, as indicated by red arrowhead. Sup., supernatant. For aptamer pull-down assays, 5′-biotinylated aptamers (IDT) were incubated with cell lysates and captured using streptavidin magnetic beads (Tamagawa Seiki Co., Ltd.). After washing, bound proteins were eluted and analyzed by western blotting using anti-CD44 antibody. HepG2 lysate was used as a control lysate for assay validation.

**Supplementary Figure 3.**
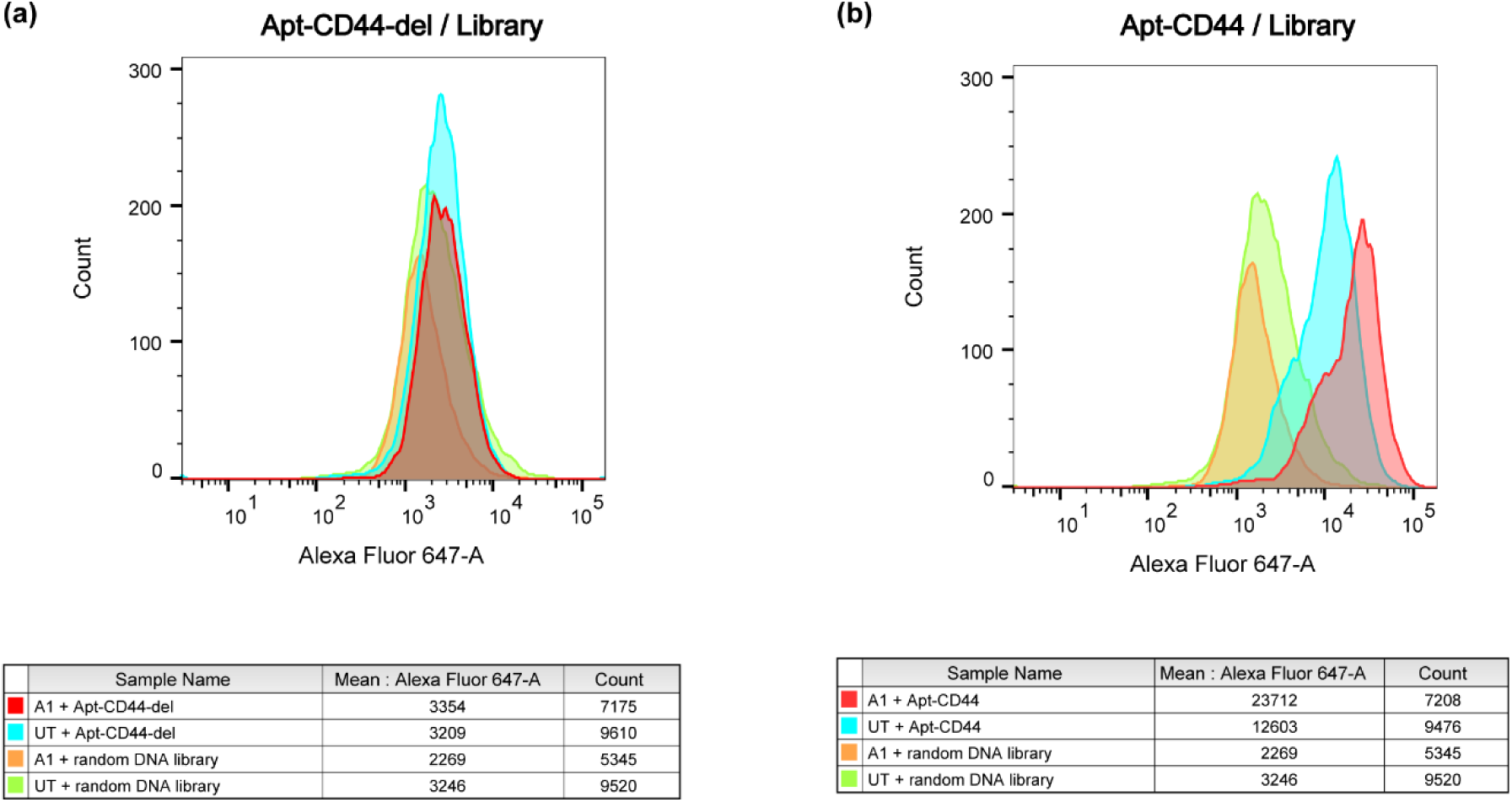
Representative flow cytometry histograms of aptamer staining. Representative histograms showing Alexa Fluor 647 fluorescence intensity in untreated (UT) and A1-stimulated astrocytes stained with (a) Apt-CD44-del control aptamer or (b) Apt-CD44. Random DNA library-stained samples are shown as additional controls. Mean fluorescence intensity and event counts for the gated single-cell population are shown below each histogram

**Supplementary Figure 4.**
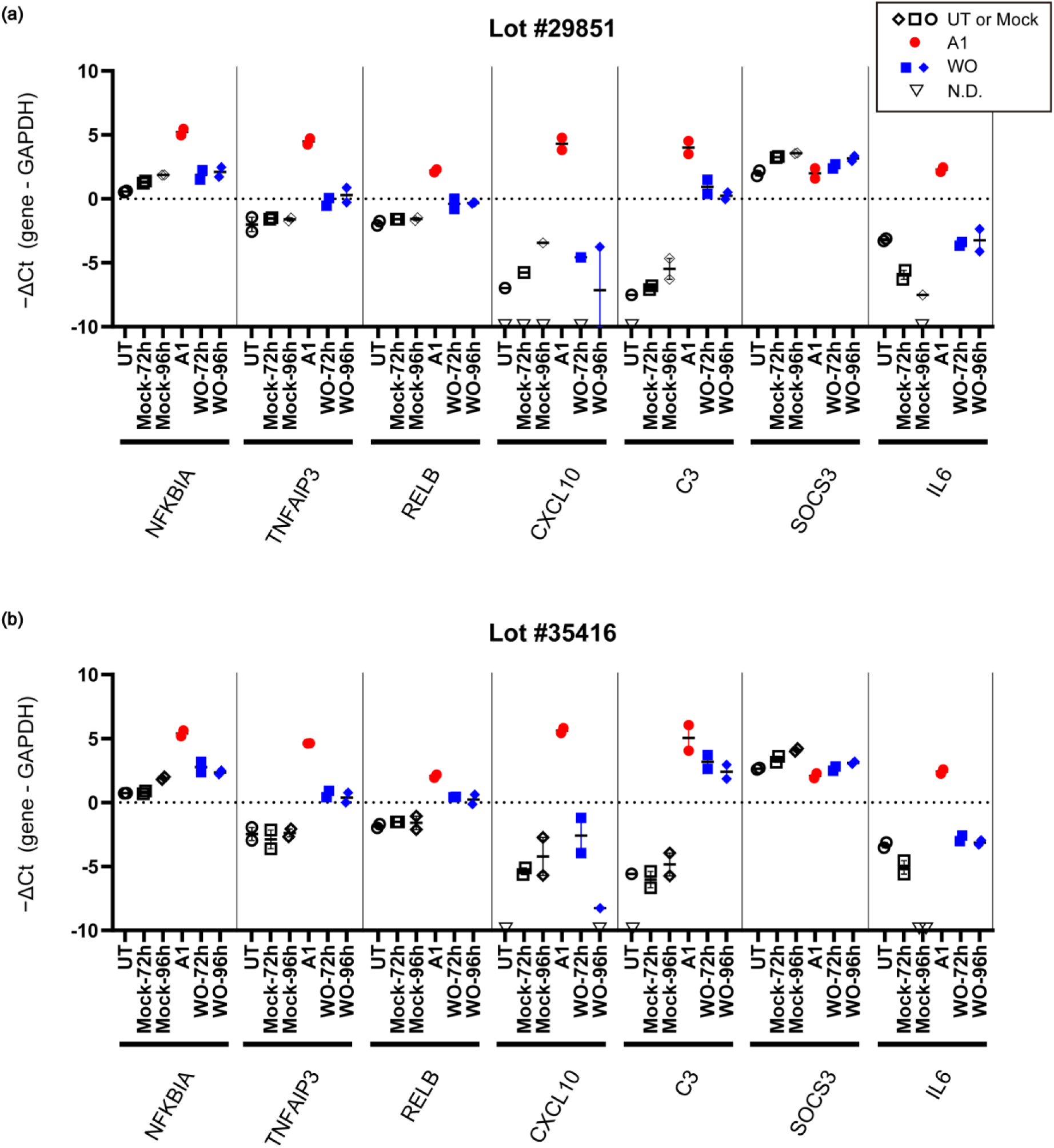
Sustained C3 expression after A1 stimulus withdrawal in an independent astrocyte lots. GAPDH-normalized expression of selected genes in untreated (UT) astrocytes, mock-treated astrocytes at 72 h and 96 h, and post-washout (WO) astrocytes at 72 h and 96 h after A1 stimulus withdrawal. Values are shown as −ΔCt. Non-detected samples are indicated as N.D.

**Supplementary Figure 5.**
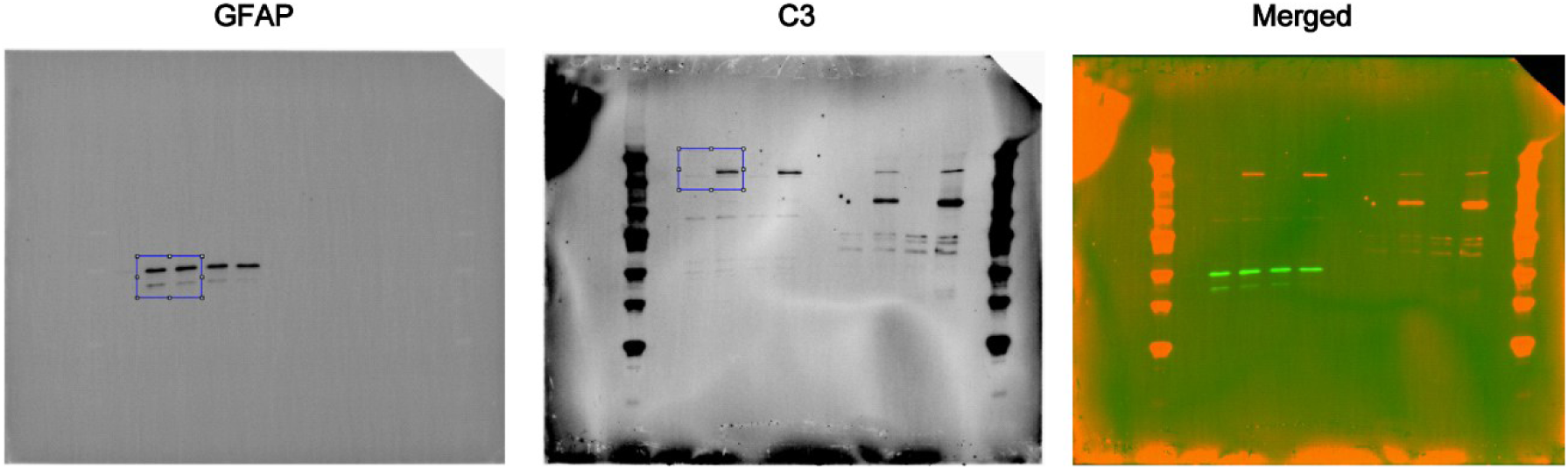
Uncropped western blot images corresponding to Fig. 1g. The regions used in the main figures are indicated by boxes.

**Supplementary Figure 6.**
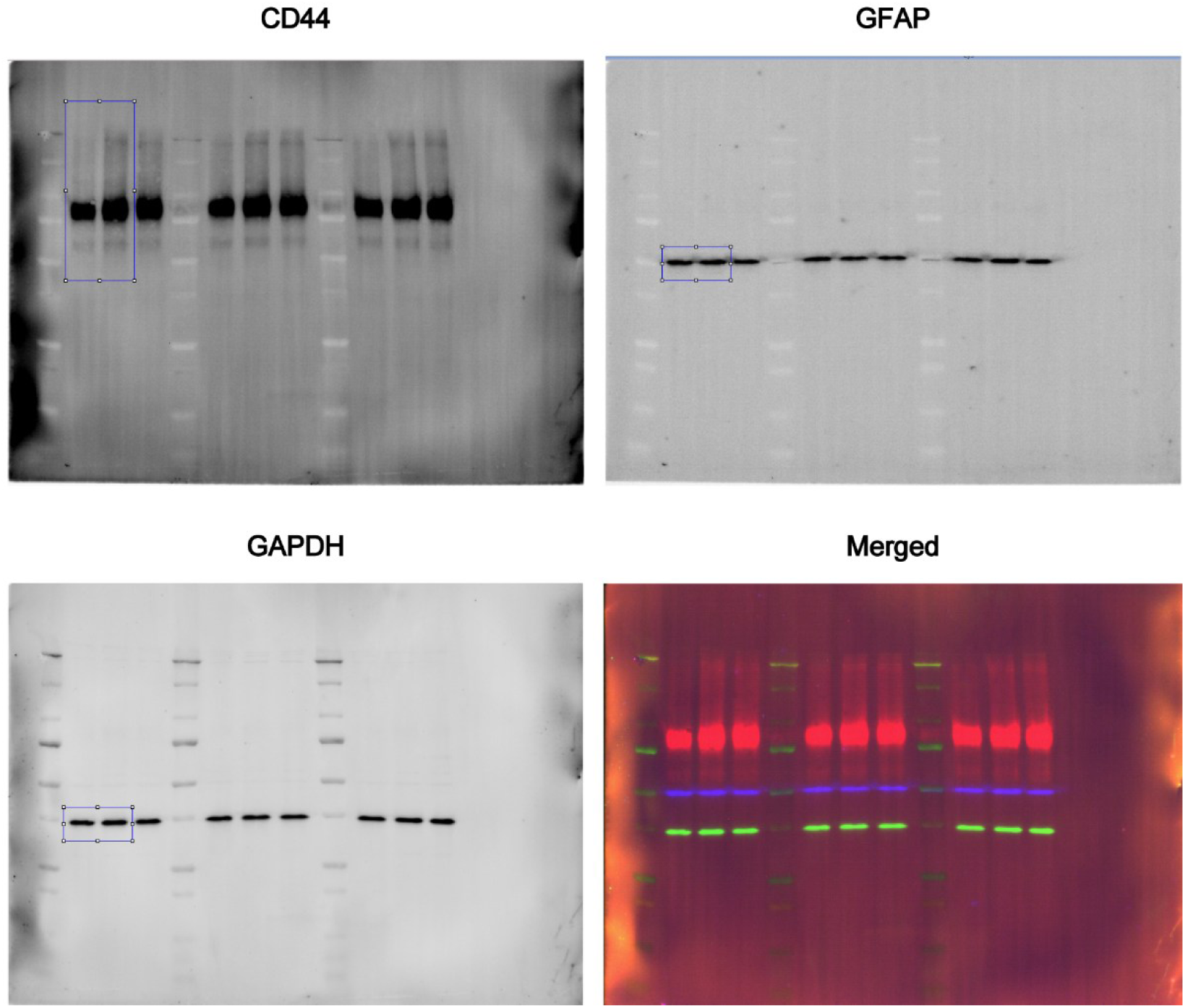
Uncropped western blot images corresponding to Fig. 2b. The regions used in the main figures are indicated by boxes.

**Supplementary Figure 7.**
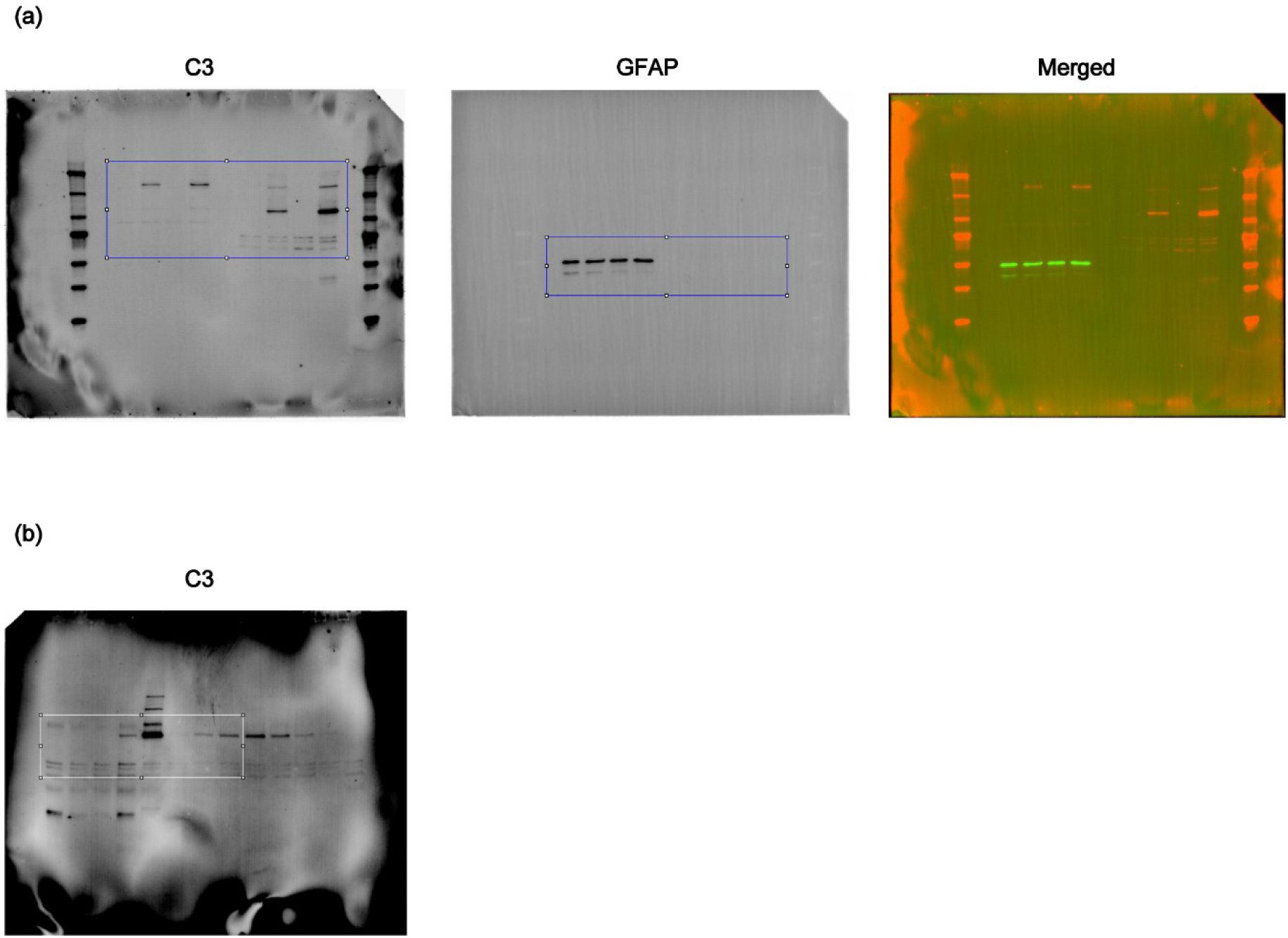
Uncropped western blot images corresponding to (a) Fig. 3f and (b) 3g. The regions used in the main figures are indicated by boxes.

